# Epigenetic regulation of epithelial dectin-1 through an IL-33-STAT3 axis in allergic disease

**DOI:** 10.1101/2020.11.01.364067

**Authors:** Hwan Mee Yong, Naina Gour, Deepika Sharma, Syed Muaz Khalil, Andrew P. Lane, Stephane Lajoie

**Author notes:** Correspondence to: Stephane Lajoie, Ph.D., Department of Otolaryngology, Johns Hopkins School of Medicine, Baltimore. These authors contributed equally (Hwan Mee Yong, Naina Gour). Thermo Fischer Scientific, Hunt Valley, MD 21031.

## Abstract

Allergic diseases arise in susceptible individuals in part because of decrements in protective pathways. The mechanism by which these anti-inflammatory molecules become repressed remains unclear. We have previously reported that epithelial dectin-1 prevents aberrant type 2 responses and is downregulated in the epithelium of allergic patients. Here we report that dectin-1 is constitutively expressed by the respiratory epithelium in humans, and that IL-33 specifically acts as a repressor of dectin-1. Mechanistically, this occurs via IL-33-dependent STAT3 activation and the subsequent repression of the dectin-1 gene, CLEC7A. We have identified a novel enhancer region upstream of the proximal promoter of CLEC7A that is only accessible in epithelial cells, but not in hematopoietic cells. Epigenetic repression of CLEC7A through this newly identified locus, downstream of an aberrant IL-33-STAT3 axis, occurs in the epithelium of allergic individuals. Collectively, our data identify a mechanism of epigenetic fine-tuning of dectin-1 expression in epithelial cells that may participate in allergenicity.

## Introduction

Allergic diseases such as asthma, allergic rhinitis, chronic rhinosinusitis with nasal polyps (CRSwNP) are driven by dysregulated type 2 responses. There are several pathways that prevent overzealous activation of these responses and confer protection to the development of allergy. In susceptible individuals a decrement in these protective mechanisms can drive disease upon exposure to sources of allergens^1-3^. Several molecular pathways, including pattern recognition receptors like CLEC10A, TLR2, TLR7/8, and TLR9, as well as well-known anti-inflammatory intracellular mediators like A20, have been shown to be protective in asthma^2-5^. Some of these pathways are epithellialy-expressed and their activity can be decreased in allergic patients^2,6^. Consistent with this, we observed lower levels of dectin-1, a c-type lectin receptor (CLR) encoded by the *CLEC7A* gene, in epithelial cells from individuals with asthma and CRSwNP and found that it protects against the development of allergic asthma by recognizing invertebrate tropomyosin in house dust mite (HDM) extract and by preventing the aberrant release of IL-33, an epithelial-derived type 2-promoting cytokine^1^.

It is thought that in susceptible individuals, a combination of genetic and epigenetic mechanisms contributes to decrements in protective pathways. Single nucleotide polymorphisms (SNP) have been associated with the regulation of asthma-protective genes. For example, a coding SNP in the A20 gene, TNFAIP3, impairs protein function and increases the propensity of developing asthma^2^. Further, we found an intronic SNP in the *CLEC7A* locus associated with lower expression of *CLEC7A* in children with asthma^1^. However, in addition to genetic mechanisms of gene regulation, several epigenetic modifications in pro-inflammatory and anti-inflammatory genes have been identified in patients with allergic diseases^7-10^. These epigenetic mechanisms can, in combination with SNPs, control gene expression in disease, and have also been shown to regulate these transcripts in the asthmatic epithelium^10-12^. Epigenetic pathways involve not only DNA methylation and non-coding RNAs, but also histone modifications, which work in concert with repressors, enhancers and transcription factors to regulate gene expression^7,10^. The importance of these mechanisms in allergy is especially evident in studies done in T cells. For example, enhancer regions in the genes coding for type 2 cytokines in T cells are controlled by transcription factors like STAT6 and GATA3 and confer susceptibility to allergic diseases^13,14^. Moreover, a recent report shows differentially regulated enhancer regions are associated with upregulated and downregulated genes in epithelial cells from asthmatics^12^. Despite this, little remains known about the epigenetic regulation of gene expression in the epithelium and the identity of external mediators that can influence this biology.

Based on this, we set out to determine whether there are epigenetic mechanisms that repress epithelial dectin-1 expression in allergy. Understanding how these restorative mechanisms become dysregulated is an essential aspect in expanding our knowledge of allergic disease pathogenesis. Here we report that dectin-1 expression is inhibited by IL-33 in human and mouse respiratory epithelial cells, but not in immune cells. Our findings show the existence of a previously unknown enhancer region upstream of the dectin-1 gene that confers unique regulation of dectin-1 in the epithelium. We find that this region of chromatin can be bound by IL-33-induced STAT3 and acts to repress dectin-1 gene expression.

## Materials and Methods

### Mice

BALB/c mice were obtained from Jackson Labs (Bar Harbor, Maine) and bred in our facility. *Il33*^*cit/cit*^ mice on the BALB/c background (gift from Dr. Andrew McKenzie) were bred to BALB/c to generate heterozygous animals. *Il33*^*cit/+*^ mice were used to generate homozygous progeny that was used for experiments. Mice were housed in a specific pathogen free animal facility in micro-isolator cages. Mice were provided autoclaved food (Lab diet 5010) and water ad libitum. All procedures were approved by the Animal Care and Use Committee of Johns Hopkins University.

### House dust mite (HDM)

HDM was obtained from Stallergenes Greer. Vials used in experiments had Der p 1 content ranging from 114.21-245.95 microgram/vial, and endotoxin 42.25-2874 EU/vial.

### HDM *in vivo* administration

*Il33*^*+/+*^ and *Il33*^*cit/cit*^ mice were given PBS (40 ul) or 50 ug HDM intratracheally on days 0 and 14. Mice were sacrificed on day 17 for analysis of dectin-1 expression.

### Flow cytometry

Mouse lung cells were obtained by digestion of lung tissue with 0.05 mg/ml Liberase TL (Roche) and 0.5 mg/ml DNaseI (Sigma) for 45 min at 37**°**C in 5% CO_2_. Digested tissue was filtered through a 70-um nylon mesh (BD Biosciences) and centrifuged. Pellet was resuspended in red blood cell lysis buffer (ACK lysis buffer). Recovered cells were counted (trypan blue exclusion), plated at 4-5×10^6^ cells/ml. Cells were washed with PBS and labeled with Zombie Aqua live/dead dye (Biolegend) for 10 min at RT, and blocked with 10 ug/ml anti-CD16/32 (BioLegend or BioXCell) for an additional 20 min at RT. Cells were stained with fluorochrome-labeled antibodies as detailed in Table 1. Human sinonasal tissue was processed similarly to mouse lungs. Cells were labeled with live/dead dye (Zombie Aqua) for 10 min at RT and blocked with human TruStain FcX (BioLegend) for 20 min at RT. See Table 1 for a list of antibodies used. Data was acquired on an LSRII flow cytometer (BD Biosciences) and gated to exclude debris and to select single cells (SSC-W/SSC-A). Data was analyzed using FlowJo (BD Biosciences).

### Human bronchial epithelial cell line culture

16HBE14o- (Sigma) and BEAS-2B (ATCC) were maintained in DMEM media (10% FBS). For transfections, 16HBE were seeded at 10,000 cells/well of a 96-well flat-bottom dish and transfected using FuGENE HD (Promega).

### SNEC and NHBE cell culture

Human nasal epithelial cells and sinonasal mucosa were obtained from nasal brushings and tissue removed during endoscopic sinus surgery from control and patients with or without nasal polyps as previously described. The research protocol was approved through the Johns Hopkins Institutional Review process, and all subjects gave signed informed consent. Inclusion criteria included continuous symptoms of rhinosinusitis as defined by the AAO-HNS Chronic Rhinosinusitis Task Force for greater than 12 weeks, computed tomography of the sinuses revealing isolated or diffuse sinus mucosal thickening or air-fluid levels, and nasal polyps visible on diagnostic endoscopy. None of the subjects had a history of tobacco use, cystic fibrosis, ciliary dyskinesia, systemic inflammatory or autoimmune disease, or immunodeficiency. Characteristics of the subjects are summarized in Table 2. Isolated SNECs cells were grown in Bronchial Epithelial cell Growth Media (BEGM bullet kit, Lonza). At 80% confluency, cells were lifted using trypsin-EDTA (0.05%, Gibco) and seeded into a 96-well culture plate pre-coated with fibronectin-collagen. NHBE (Lonza) were cultured similarly to SNECs. Upon reaching 80-90% confluency, cells were starved overnight in BEGM media without the bovine pituitary extract (BEGMnoBP). Cells were treated with HDM (100 ug/ml) for 2h and cells were harvested for RNA. Where indicated, cells were pretreated for 2h with 5 uM stattic (Cayman) or DMSO, or 5 ug/ml isotype or neutralizing monoclonal anti-hIL-33 antibodies (R&D Systems, MAB36254), followed by stimulation with rIL-33 (Peprotech), HDM (100 ug/ml) for 2h. RNA was extracted (TRIzol), and reverse-transcribed to analyze *CLEC7A* expression. Primers are designed to span an intron to avoid co-amplification of genomic DNA (Table 3).

### Phosphoflow

CRSwNP hSNECS were cultured in collagen/fibronectin-coated 6-well plates in BEGM media. Upon reaching 80-90% confluency, cells were starved overnight in BEGMnoBP. Cells were stimulated for 2h with media or recombinant human IL-33 (Peprotech). Cells were then placed on ice, washed with ice-cold PBS and incubated with ice-cold 0.25% trypsin on ice for 25-35 mins. Cells were detached with gentle pipetting and 32% paraformaldehyde (Electron Miscroscopy Sciences) was directly added to the media for a final concentration of 1.6%, and fixed for 10 min at RT. Cells were transferred to 15 ml conical tubes, spun, PFA was removed and cells were permeabilized with 500 µl ice-cold 90% methanol for 60 min at 4 °C. Cells were washed once in PBS+0.2% BSA and stained for 30 min with anti-human STAT3 pTyr705-Alexa Fluor 647 (BioLegend) in PBS+0.2% BSA at RT. Cells were washed twice in PBS+0.2% BSA.

### Transfection of SNECs and NHBE

Transfection was done in BEGM media in 96-well dishes (65-85% confluent) with 0.098 ug *CLEC7A*.luc and 0.002 ug pGL4.PGK control vector per well (Promega) using Trans-IT X2 (Mirus). The following day, cells were starved in BEGMnoBP overnight. Media was removed, cells were rinsed once in PBS, and 150 ul/well BEGMnoBP was added, after 4-6h, 25 ul was collected for determination of secreted Nano-Glo, and cells lysed (25 ul/well, Passive Lysis Buffer, Promega) for detection of the firefly control (pGL4-PGK).

### Mouse tracheal epithelial cells

Mouse tracheal epithelial cells were isolated from tracheas digested overnight at 4 °C in Ham’s F12 medium plus pronase (1 mg/ml; Roche). Cells were cultured for 3 h on Primaria plates (Falcon) for removal of fibroblasts. Non-adherent cells were resuspended in BEGM and plated at 25,000 cells per well of a 96-well dish. The following day, cells were harvested for RNA and determination of Clec7a expression (Table 3 for primers).

### Dendritic cell culture

Human normal dendritic cells (Lonza) were cultured in 50 ng/ml GM-CSF and 50 ng/ml IL-4 for 5 days, after which they were stimulated with HDM (100 ug/ml) in combination with 10 ug/ml isotype control or neutralizing anti-IL-33 antibodies for 4h.

### Plasmids

To generate *CLEC7A*.luc reporter vectors, a 3.5kb fragment of the human *CLEC7A* promoter region (chr12:10129421-10132905) was synthesized (GenScript) and cloned into the secreted Nano-Glo luciferase vector pNL1.3 (Promega). Mutation of the STAT3 binding site (wt: TTTCCAAGCAC, mut: TGGAAGGGCAC) in *CENSER* was performed commercially (GenScript). CENSER and proximal deletion constructs were done using the QuikChange mutagenesis kit (Agilent). Deletions were verified by gel electrophoresis.

### Chromatin immunoprecipitation

0.5-1.0×10^6^ epithelial cells (BEAS-2B) were trypisinized and fixed in 1% PFA (8 min) followed by quenching with 1 M glycine (10 min). Cells were washed twice in ice-cold PBS before storage at −80°C. Cells were lysed in 150 uL lysis buffer (1% SDS, 50 mM Tris-Cl (pH 8.1), 10 mM EDTA (pH 8.0)) and sonicated in TPX microtubes (Diagenode) in a 4°C-cooled water bath sonicator (Diagenode Bioruptor, HI setting, 30 sec ON/30 sec OFF) for 25-30 cycles to properly shear the DNA. Lysates were cleared by centrifugation (10 min, 18,000xg) and pre-cleared for non-specific binding with 40 ul BSA-blocked protein A/G magnetic Dynabeads (Invitrogen) for 4h at 4°C. Lysates were diluted 10-fold (1.1% Triton X-100, 0.01% SDS, 167 mM NaCl, 16.7 mM Tris-Cl pH 8.1, 1.2 mM EDTA) and incubated with 3.5 ug anti-STAT3 (clone ST3-5G7, Invitrogen) or anti-GFP (Santa-Cruz) antibodies overnight at 4°C. Antibodies were captured with 40 ul BSA-blocked protein A/G magnetic Dynabeads on a rocking platform for 4h at 4°C. Beads were washed in 1 mL low salt buffer (0.1% SDS, 1% Triton X-100, 2 mM EDTA, 20 mM Tris pH 8.1, 150 mM NaCl), then, high salt buffer (0.1% SDS, 1% Triton X-100, 2 mM EDTA, 20 mM Tris pH 8.1, 500 mM NaCl), lithium chloride buffer (0.25 M LiCl, 1% Nonidet P-40, 1% deoxycholate, 1 mM EDTA, 10 mM Tris pH 8.1), followed by 2 washes in Tris-EDTA (pH 8.1). Chromatin was eluted from the beads using 200 ul elution buffer for 15 min at RT (1% SDS, 0.1 M NaHCO_3_), followed by de-crosslinking in 0.3 M NaCl at 65°C overnight. Chromatin was RNase-digested (30 min at 37°C), followed by proteinase K (1h at 56°C), and purified using PCR purification columns (Qiagen). PCR was performed using primers in the *CLEC7A* locus (see Table 3). All data was normalized to input and presented as enrichment over the *IGX1A* negative control region (primers from Qiagen).

## Results

### Respiratory epithelial cells express dectin-1

While the expression, regulation and function of prototypical PRRs such as TLRs have been extensively studied in the allergic epithelium, our understanding of the role of CLRs in allergic diseases is mostly unclear and has majorly been derived from their expression on hematopoietic cells (macrophages, dendritic cells, etc). Exploration of CLRs in human tracheal and bronchial epithelial cells (RNA-seq dataset GSE101993) reveals that several are constitutively expressed, including CLEC7A, the gene coding for dectin-1, suggesting that they may play an important role in regulating epithelial function (Fig. 1a). The majority of these CLRs are poorly understood, and many have unknown function, especially on the epithelium. Dectin-1 is probably the best studied CLR and its function is relatively well understood in myeloid cells in context of fungal infections. However, we^1^ and others^15-20^ have reported expression of dectin-1 in epithelial cells, and found that it has a key role in preventing aberrant IL-33 release in response to dust mite allergen. Further analysis of the CLEC7A locus using ENCODE data shows the transcribed exons across the human respiratory epithelium in nasal, tracheal, bronchial and airway epithelial cells (Fig. 1b). These data support the notion of epithelial cells as important non-hematopoietic dectin-1-expressing cells. In humans and mice, respiratory epithelial cells can be divided into subsets that can be deconvoluted based on EpCAM and CD104 or CD49f expression^21-24^. Where EpCAM^+^CD104^-^ are luminal cells, EpCAM^+^CD104^+^ are luminal progenitors and EpCAM^lo^CD104^+^ represent basal cells. We sought to characterize the expression of dectin-1 in these subsets. To this end, we used human sinonasal tissue, and we found that luminal progenitor and basal epithelial cells have the highest levels of dectin-1-expressing cells, with marginal expression in luminal epithelial cells (Fig. 1c). Our findings demonstrate that the epithelium maintains tonic levels of dectin-1 across the respiratory system.

**Figure 1.**
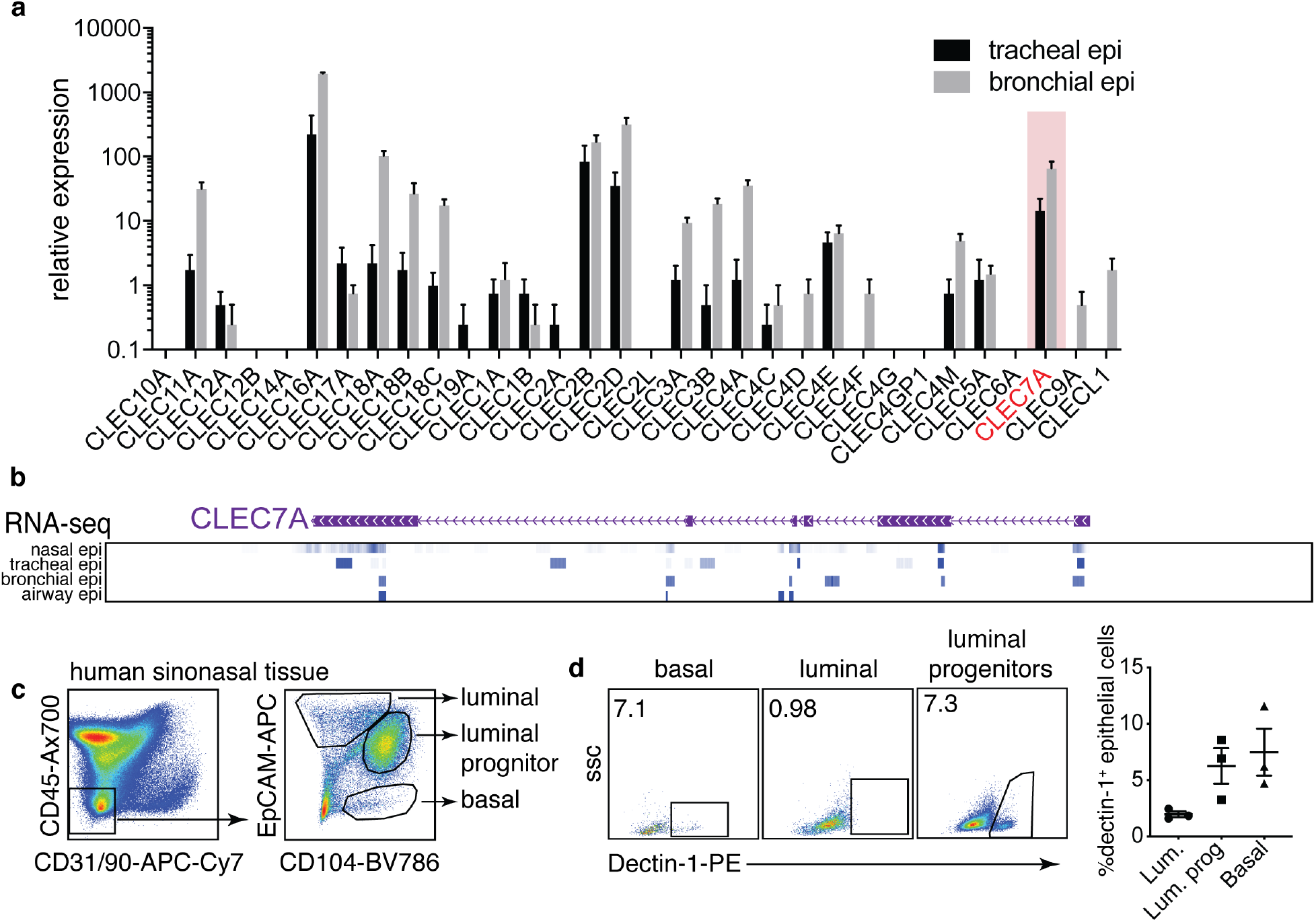
Respiratory epithelial cells express dectin-1. (a) Levels of CLR mRNAs in human bronchial and tracheal epithelial cells from healthy donors (RNA-seq, GSE101993). (b) RNA-seq from ENCODE highlighting CLEC7A exon transcription in nasal, tracheal, bronchial and airway epithelial cells. (c) gating scheme identifying various lung epithelial subsets by flow cytometry and (d) expression of dectin-1 in these populations. Data is means+SEM and pooled from 4 (a) or 3 (d) donors.

### Allergen-induced IL-33 represses epithelial dectin-1

Our previous work has shown that dectin-1 is downregulated in the allergic epithelium, and that it functions to inhibits allergen-induced IL-33^1^. Based on this, we postulated that dectin-1 is repressed by an autocrine pathway. As IL-33 is enriched in allergic airway tissue, we hypothesized that epithelial IL-33 may counter dectin-1 expression. To begin to explore this, we treated cultured human sinonasal epithelial cells (hSNECs) from nasal polyp patients with media or HDM and treated with isotype control or blocking anti-IL-33 mAbs. Interestingly, we observed that HDM reduced *CLEC7A* expression, and this repression was restored when IL-33 was neutralized (Fig. 2a). Interestingly, these culture conditions had no effect on expression of *CLEC7A* in human primary dendritic cells (Fig. 2b), suggesting a cell-type specific regulation of dectin-1. In line with this, treating hSNECs with rIL-33 (250 pg/ml) inhibited *CLEC7A* expression (Fig. 2c). Consistent with our human data, isolated mouse tracheal epithelial cells from IL-33 deficient (*Il33*^*cit/cit*^) mice express more *Clec7a* than wildtype cells (Fig. 2d). Next, we explored the possibility that exposure to HDM could enhance responsiveness to autocrine IL-33 via upregulating its receptor, IL1RL1/ST2. We found that treating CRSwNP SNECs with HDM upregulated *IL1RL1* (ST2) mRNA, but not in controls (supplementary Fig. 1) and suggests that allergen exposure in allergic epithelial cells enhances their production and responsiveness to IL-33. Overall, these data suggest a functional pathway in mice and human in which IL-33 represses epithelial dectin-1 expression.

**Figure 2.**
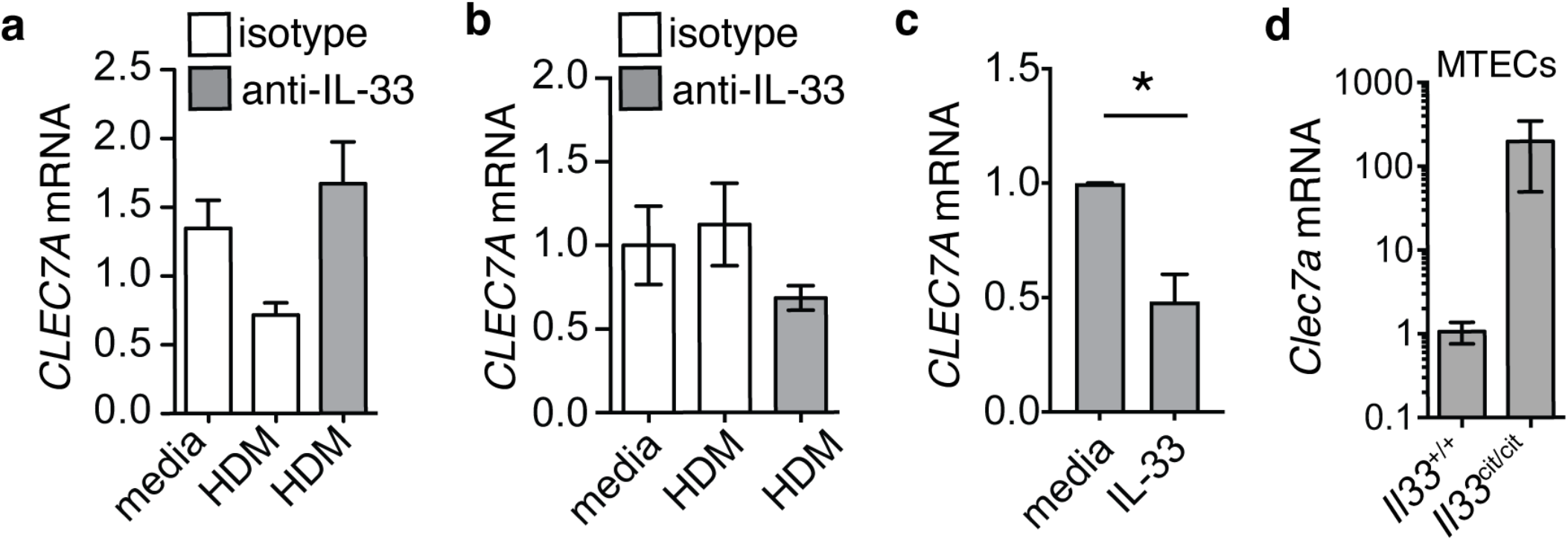
Allergen-induced IL-33 represses epithelial dectin-1. CLEC7A mRNA levels in (a) CRSwNP SNECs or (b) dendritic cells cultured in the presence of media, HDM (100 ug/ml) in combination with isotype or blocking anti-IL-33 antibodies (5 ug/ml) for 2h. (c) CLEC7A mRNA levels in CRSwNP SNECs treated with media or 0.25 ng/ml IL-33 for 2h. (d) Clec7a mRNA in cultured mouse tracheal epithelial cells from wildtype or *Il33*^*cit/cit*^ mice. Data is means+SEM and represents representative data from 3 donors or 3 independent experiments.

### IL-33 represses epithelial dectin-1 *in vivo*

To further examine the regulation by IL-33 of dectin-1 *in vivo*, we exposed, previously characterized IL-33 deficient (*Il33*^*cit/cit*^) mice^25^, to PBS or HDM. We find that lack of IL-33 results in an increase in the percentage of dectin-1^+^ basal epithelial cells in mice treated with HDM, when compared to control (Fig. 3a,b). Similarly, we find that the frequency of dectin-1^+^ luminal progenitor epithelial cells increases significantly in *Il33*-deficient animals after HDM exposure, as compared to IL33-sufficient mice (Fig. 3c,d). Surprisingly, we show that, while dectin-1 expression is abundant in myeloid cells, IL-33 does not appear to regulate it. Various myeloid populations were identified (see gating approach, supplementary Fig. 2), and we found that the frequency of dectin-1^+^ alveolar macrophages (Fig. 3e), interstitial macrophages (Fig. 3f), and CD11b^+^ dendritic cells (Fig. 3g) remains unaffected in *Il33*^*cit/cit*^ animals. Overall, our data suggest that IL-33 represses the expression of dectin-1 specifically in epithelial cells and not hematopoietic cells, implying the involvement of cell-type specific epigenetic regulation of dectin-1. We did not observe that HDM caused a decrease in the frequency of mouse dectin-1^+^ epithelial cells, as it does in human nasal polyp-derived epithelial cells, suggesting that HDM may elicit non-epithelial factors that could promote survival or proliferation of respiratory epithelia. Nonetheless, these data overall demonstrate that allergen-induced IL-33 specifically represses epithelial dectin-1 in mouse and humans.

**Figure 3.**
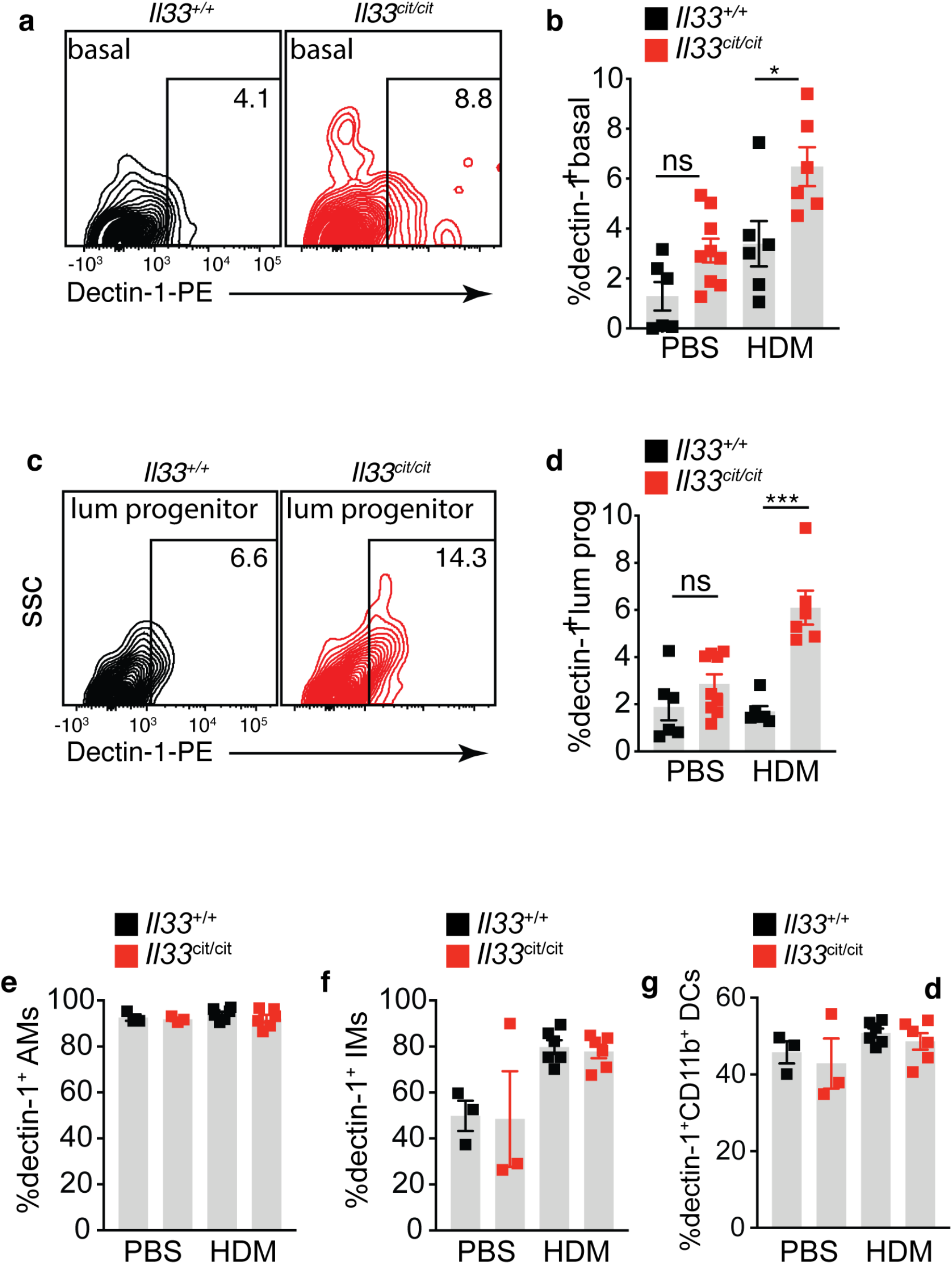
IL-33 represses epithelial dectin-1 *in vivo*. Il33^+/+^ and Il33^cit/cit^ mice were treated with PBS or HDM and lung dectin-1^+^ cells were analyzed. Dectin-1^+^ basal epithelial cells in (a) representative flow plots and (b) frequencies. Dectin-1^+^ luminal progenitors in (c) representative flow plots and (d) frequencies. Dectin-1^+^ (e) alveolar macrophages (f) interstitial macrophages and (g) CD11b^+^ dendritic cells. Data is means+SEM and representative of at least 2 independent experiments. Data was analyzed by one-way ANOVA followed by post-hoc test. *p<0.05; ***p<0.001.

### Identification of a functional epithelial-specific enhancer region in the *CLEC7A* loci

To understand the epithelial-specific regulation of dectin-1, we investigated its regulatory landscape using ENCODE data where we analyzed DNaseI hypersensitive sites (DHS-seq) from primary human epithelial cells from various tissues, as well as from hematopoietic cells such as CD14^+^ monocytes and CD1c^+^ dendritic cells. This analysis revealed a previously reported proximal promoter^26^, which is active in both cell types, as demonstrated by an open chromatin configuration (Fig. 4a). In addition, we found a second, previously unreported, distal site of open chromatin, seen only in epithelial cells, we refer to this region as the ClEc7a Nonhematopoietic-SpEcific Region-CENSER. This suggests that CENSER acts as a unique regulatory region, differentially regulating dectin-1 expression between non-hematopoietic and hematopoietic cells. Moreover, we found this region to be enriched for the histone modifications H3K4me1 and H3K4me3 in epithelial cells (Supplementay Fig. 3), further supporting the role of CENSER as a regulator of CLEC7A expression in the epithelium. To evaluate the function of these regulatory regions of chromatin in controlling CLEC7A expression, we generated a 3.5kb promoter construct containing both the proximal and CENSER region upstream of secreted NanoLuc. We observed significant activity of this CLEC7A reporter construct in transiently transfected primary human sinonasal epithelial cells (Fig. 4b). Deletion of the proximal region attenuated *CLEC7A* promoter activity in primary human nasal epithelial cells, consistent with the important role of proximal promoters in regulating gene function (Figure 4b). Notably, removal of CENSER also reduced *CLEC7A* promoter activity in SNECs (Fig. 4b). Similarly, deletion of the CENSER and proximal regions also decreased *CLEC7A* promoter activity in human bronchial epithelial (NHBE) cells (Fig. 4c). Overall our data suggests a role for this CENSER region as an enhancer regulating dectin-1 expression in epithelial cells.

**Figure 4.**
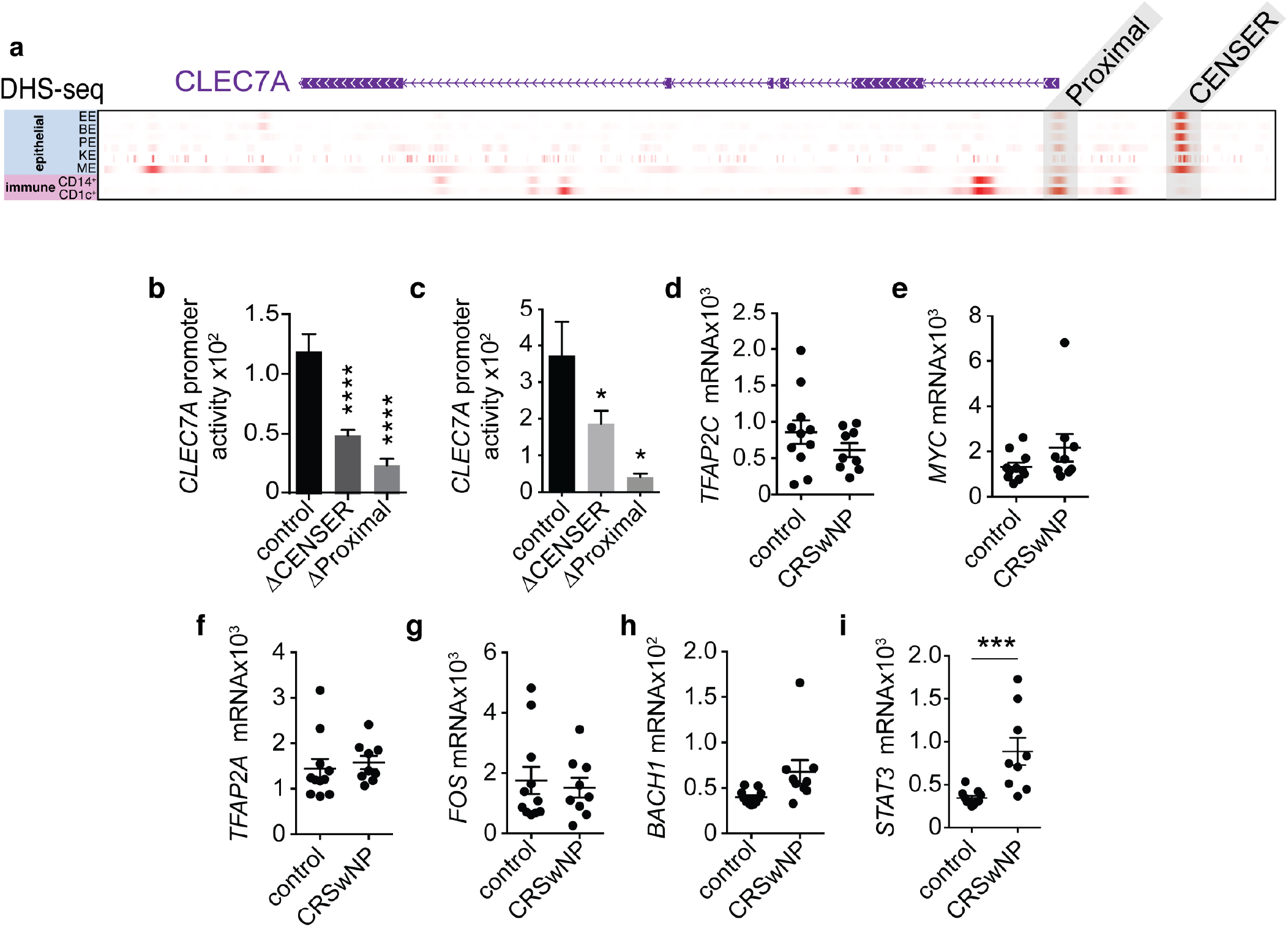
Identification of a functional epithelial-specific enhancer region in the *CLEC7A* loci. (a) DHS-seq data from ENCODE for the CLEC7A locus for primary esophageal (EE), bronchial (BE), prostate (PE), and mammary epithelial (ME) cells, keratinocytes (KE), CD14^+^ monocytes and CD1c^+^ dendritic cells. CLEC7A promoter activity in (b) SNECs and (c) NHBE transfected with full-length or truncated promoter constructs. Levels of (d) TFAP2C, (e) MYC, (f) TFAP2A, (g) FOS, (h) BACH1, and (i) STAT3 mRNA in control and CRSwNP SNECs. Data is means+SEM and representative of 2-3 independent experiments. Data was analyzed by one-way ANOVA, Student’s T test, or Mann Whitney test (control vs. CRSwNP). *p<0.05, ***p<0.001, ****p<0.0001.

In order to elucidate the mechanism through which CENSER region controls dectin-1 promoter activity we analyzed ENCODE ChIP-seq data, and this revealed this region to be a target of the transcription factors TFAP2A, TFAP2C, MYC, FOS, BACH1 and STAT3 (Supplementary Fig. 4). We next assessed the levels of these CENSER-binding transcription factors (TFs) in cultured SNECs from control and CRSwNP patients (Fig. 4d-i). We found STAT3 mRNA to be significantly higher in the allergic epithelium (Fig. 4i). Consistently, exposure to HDM has been shown to activate STAT3 in mouse lung epithelial cells, leading to allergen-induced experimental asthma^27,28^. These data suggest a pro-allergic function for STAT3 where it could act as repressor of dectin-1 during aberrant Th2 responses.

### STAT3 represses CLEC7A through CENSER

IL-33 has been shown to phosphorylate STAT3 in skin, gut and breast epithelial cells^29-32^. Here we confirm that at low (250 pg/ml) or more standard (10 ng/ml) concentrations, IL-33 leads to an increase in the frequency of STAT3 pTyr705^+^ hSNECs (Fig. 5a,b). Next, to test the direct role of STAT3-CENSER interaction in controlling dectin-1 expression, we performed site-directed mutagenesis of the CENSER STAT3 binding site. We found that disrupting the STAT3 binding site significantly enhanced *CLEC7A* promoter activity in bronchial epithelial cell line (Fig. 5c), establishing the repressive function of STAT3 binding at the *CLEC7A* locus. Analysis of NCBI GEO data (GDS3106) also demonstrates that levels of *Clec7a* are higher in *Stat3*^*-/-*^ mouse tracheal epithelial cells, as compared to controls (Fig. 5d). Next, we performed ChIP-PCR in bronchial epithelial cell line to assess STAT3 binding at the CLEC7A locus in response to IL-33. Using primers to amplify the regions around or at the proximal and CENSER, we found that, compared to control (anti-GFP), IL-33 promoted recruitment of STAT3 to the CENSER region, but not others (Fig. 5e). STAT3 is not only activated by IL-33, but can act as a transcriptional repressor^33-35^. Based on that and the fact that it can be enriched at CENSER, we hypothesized that an IL-33-STAT3 axis acts to repress CLEC7A transcription. We set out to test whether inhibition of STAT3 could upregulate CLEC7A in cultured CRSwNP SNECs. We found that HDM- (Fig. 5f) and IL-33-mediated (Fig. 5g) repression of *CLEC7A* in nasal polyp epithelial cells could be completely restored by the STAT3 inhibitor stattic. Taken together, these findings reveal that an aberrant IL-33-STAT3 axis represses *CLEC7A* through its interaction with a newly described enhancer region. These data uncover a potential novel epigenetic mechanism underlying the regulation of CLEC7A in the respiratory epithelium.

**Figure 5.**
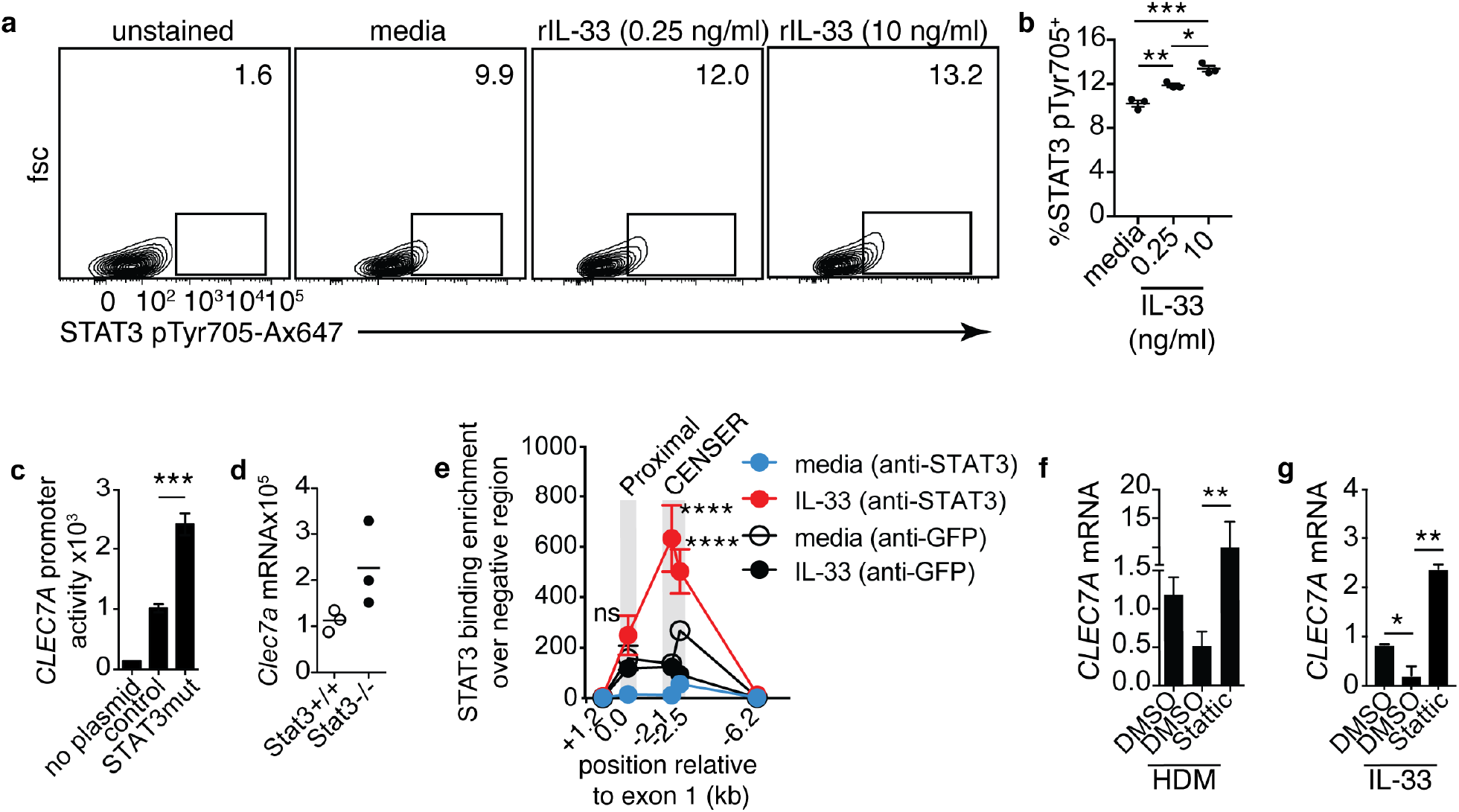
STAT3 represses CLEC7A through CENSER. (a) representative flow plots and (b) frequencies of STAT3 pTyr705 phosphoflow in CRSwNP hSNECs treated with IL-33 for 2h. CLEC7A promoter activity in (c) bronchial epithelial cell line or (d) Clec7a mRNA in isolated mouse tracheal epithelial cells from *Stat3+/+* and *Stat3-/-* mice. (e) STAT3 binding at the CLEC7A locus in bronchial epithelial cell line treated with media or IL-33 (10 ng/ml) for 2h. CLEC7A mRNA in CRSwNP SNECs treated with (f) HDM (100 ug/ml) or (g) IL-33 (0.25 ng/ml) and vehicle or 5 uM STAT3 inhibitor, stattic for 2h. Data is means+SEM and representative of 2-3 independent experiments, representative of 3 donors (a,b,f,g). Data was analyzed by one-way ANOVA followed by post-hoc test or two-way ANOVA followed by Bonferroni’s multiple comparisons test (panel e, where significances are displayed for IL-33 (anti-STAT3) vs. IL-33 (anti-GFP)). *p<0.05; **p<0.01; ***p<0.001, ****p<0.0001.

## Discussion

In susceptible individuals, allergic airway diseases like asthma and CRS are thought to develop due to the aberrant expression of pathogenic pathways. However, we now know that insufficiencies in protective mechanisms also contribute to disease. Recent reports have uncovered molecular pathways, including PRRs like dectin-1 and CLEC10A, that mediate protection against aberrant type 2 responses^1-4^ and where decrement in these pathways, many of which in the epithelium^1,2^, result in a greater propensity to develop allergy. Thus, understanding the function and regulation of these protective epithelial sensors will be important to explain how individuals develop susceptibility to allergy.

Dectin-1, while traditionally associated with expression on macrophages and dendritic cells, is, as we found, constitutively expressed by human respiratory epithelial cells, and it is also expressed by epithelial cells of other barrier surfaces like the skin, eyes, and gut^15-17,19,20,36-38^. Nevertheless, little remains known about the regulation and function of epithelial dectin-1. Reports have shown that stimulation of human corneal epithelial cells with heat-killed *Candida albicans* results in the upregulation of dectin-1^39^, and that it is required for the secretion of anti-fungal peptidoglycan recognition proteins (PGLYRP). Consistent with this, live, but not heat-killed, *Aspergillus fumigatus* conidia upregulated dectin-1 in primary human bronchial epithelial cells (HBE) in a TLR2-dependent manner^40^. Furthermore, exposure of HBE to *A. fumigatus* resulted in dectin-1-dependent cytokine release^40^. These findings are in line with the well-documented role of dectin-1 as a receptor for fungal β-glucan moieties, where this interaction triggers anti-fungal responses.

In addition to epithelial dectin-1 expression being triggered by fungus, some have shown that it can be induced by TLR2 ligand-containing bacteria. Specifically, while *Mycobacterium tuberculosis* (Mtb) and *Staphylococcus aureus* induced dectin-1 expression in the human alveolar epithelial A549 cell line^36^, TLR4-ligand containing bacteria, like *Escherichia coli*, could not activate dectin-1^36^. Functionally, siRNA-mediated knockdown of dectin-1 resulted in impaired cytokine response to Mtb and elevated Mtb colony forming units (CFUs)^36^. Others have shown that A549 cells overexpressing dectin-1 and infected with non-typable *Haemophilus influenzae* are driven to produce cytokines but this is abolished in cells expressing a non-functional dectin-1^20^. In the context of allergy, we have previously shown that in addition to these microbial ligands, the HDM protein allergen *Der p 10*, also known as invertebrate tropomyosin, is recognized by dectin-1 and this interaction prevent aberrant release of IL-33 by the epithelium^1^. Taken together, these data suggest that intact levels of epithelial dectin-1 promote anti-microbial responses and protect against allergic manifestations.

While some microbial interactions appear to drive dectin-1 expression in epithelial cells, the molecular mechanisms that regulate levels of epithelial dectin-1 are not understood. Although we have previously reported that an intronic SNP in *CLEC7A* is associated with lower gene expression in children with asthma^1^, epigenetic influences in response to inflammatory mediators also significantly contribute to gene expression. This supports the longstanding view that gene-environment interactions, especially in the epithelium, are regulators of susceptibility to allergy.

We found that exposure to dust-mite allergen led to a decrease in CLEC7A mRNA in epithelial cells from patients with nasal polyps (CRSwNP), which was restored by neutralizing IL-33. Moreover, mice lacking IL-33 also display significantly more dectin-1^+^ basal epithelial cells and dectin-1^+^ luminal progenitors. Consistent with this, IL-33 and other type 2-promoting innate epithelial cytokines like TSLP and IL-25 are known to downregulate molecules in human keratinocytes, such as filaggrin and claudin-1, that protect against atopic dermatitis^30,31,41^. Thus, IL-33, IL-25 and TLSP may promote disease by decreasing protective mucosal mechanisms in addition to directly driving type 2 inflammation.

We now know that many PRRs are expressed by epithelial cells, and reports have shown that they may function differently in the epithelium than they would on myeloid cells. For example, key studies have reported that epithelial TLR4 is a driver of type 2 responses to HDM^42,43^, whereas its expression on myeloid cells promotes neutrophilic responses to dust mite^43^. In addition to cell-type-specific function, PRR expression may also be differentially regulated in hematopoietic and non-hematopoietic compartments. Our data shows that while IL-33 represses epithelial dectin-1, blockade of IL-33 had no impact on haematopoietically expressed dectin-1.

We found that the cell-type specific (dys)regulation of *CLEC7A* in the human epithelium could be associated with *CENSER*, an upstream enhancer of the *CLEC7A* locus, that is in an open chromatin configuration only in epithelial cells, and not in human macrophages or DCs. Thus suggesting that this particular enhancer region confers an added level of regulation for dectin-1 expression in the epithelium. Regulatory regions that are designed to control gene expression in immune versus non-immune cells have been reported before. *An* epithelial-specific repressor in the *TLR4* locus has been found to maintain low levels of TLR4 in gut epithelial cells, while having no effect in monocytes^44^. Perhaps there is an evolutionary advantage to epigenetically regulate the expression of PRRs differentially in mucosal epithelial cells versus immune cells. Maintaining expression of protective PRRs, like dectin-1, while keeping pro-Th2 PRRs like TLR4 at low levels at the epithelial interface would favor homeostatic responses to environmental exposures.

*CENSER*, acting as an enhancer, is bound by several experimentally verified transcription factors, including STAT3. Moreover, STAT3 is significantly upregulated in nasal polyp epithelial cells as compared to controls, suggesting that STAT3 may promote type 2 responses. This is supported by other reports showing that epithelial STAT3 promotes type 2 inflammation in the lungs in response to HDM in mice^27^, and that pharmacological blockade of STAT3 inhibits both IL-13 and IL-17A responses to HDM^28^. As STAT3 has been reported to act as a transcriptional repressor^33-35^, in addition to several reports demonstrating that IL-33 activates STAT3^31,45,46^, we postulated a role for an IL-33-STAT3 axis in inhibiting dectin-1 expression in epithelial cells. Our data shows that impairing STAT3 function enhances dectin-1 gene expression, and thus establishes STAT3 as repressor of dectin-1. Moreover, treating an epithelial cell line with IL-33 leads to the accumulation of STAT3 binding at CENSER, but not at the proximal promoter region. This is similar to other reports demonstrating that repression of filaggrin and claudin-1 in atopic dermatitis is driven by an IL-33-STAT3 axis^30,31^. This supports the notion that dysregulation of STAT3 in the allergic epithelium is a mechanism that disrupts homeostatic levels of protective molecules like dectin-1. Nevertheless, while we have identified STAT3 as inhibiting the CENSER enhancer, it remains to be determined which TF acts as a driver of CLEC7A transcription through this region.

Taken together, our findings suggest the existence of a pathophysiologically relevant pathway that regulates epithelial dectin-1 expression. We propose that aberrant production of IL-33 by respiratory epithelial cells from allergic individuals results in an autocrine-mediated increase in STAT3 activation. A dysregulated IL-33-STAT3 axis then targets the *CENSER* enhancer, only accessible in epithelial cells, to repress dectin-1 gene expression. Our results provide new insights into the mechanisms by which dysregulated innate epithelial cytokines may drive allergy through the reduction of protective pathways.

## Author contributions

HMY and NG performed the experiments with help from DS and SMK. APL provided samples from control and CRS patients. NG and SL designed the experiments and performed data analysis. SL performed the *in silico* (ChIP-seq, DHS-seq, RNA-seq) analysis and wrote the manuscript.

## Acknowledgements

Hwan Mee Yong has nothing to disclose, Dr. Gour has nothing to disclose, Dr. Sharma has nothing to disclose, Dr. Khalil has nothing to disclose, Dr. Lane reports grants from NIH (R01AI132590) as support during the conduct of the study. This work was funded by the National Institute of Allergy and Infectious Diseases (R01AI127644) to S.Lajoie

## Figure legends

**Supplementary Figure 1.**
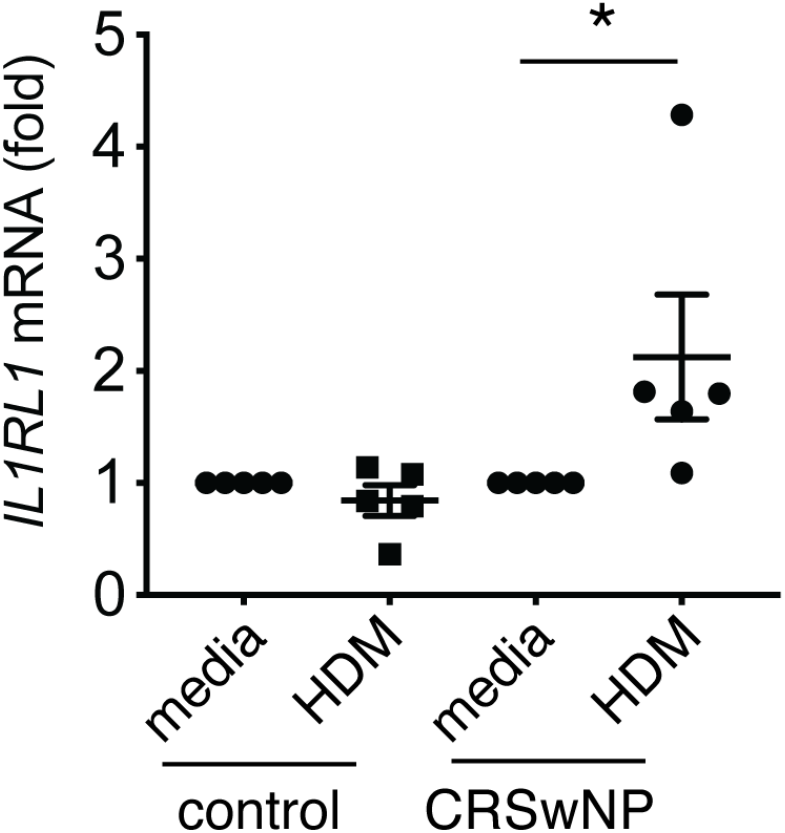
*IL1RL1* mRNA expressed as fold over media in hSNECs from control or CRSwNP patients treated with media or HDM (100 ug/ml) for 2h. Each dot represents a donor. Data was analyzed by one-way ANOVA followed by post-hoc test. *p<0.05

**Supplementary Figure 2.**
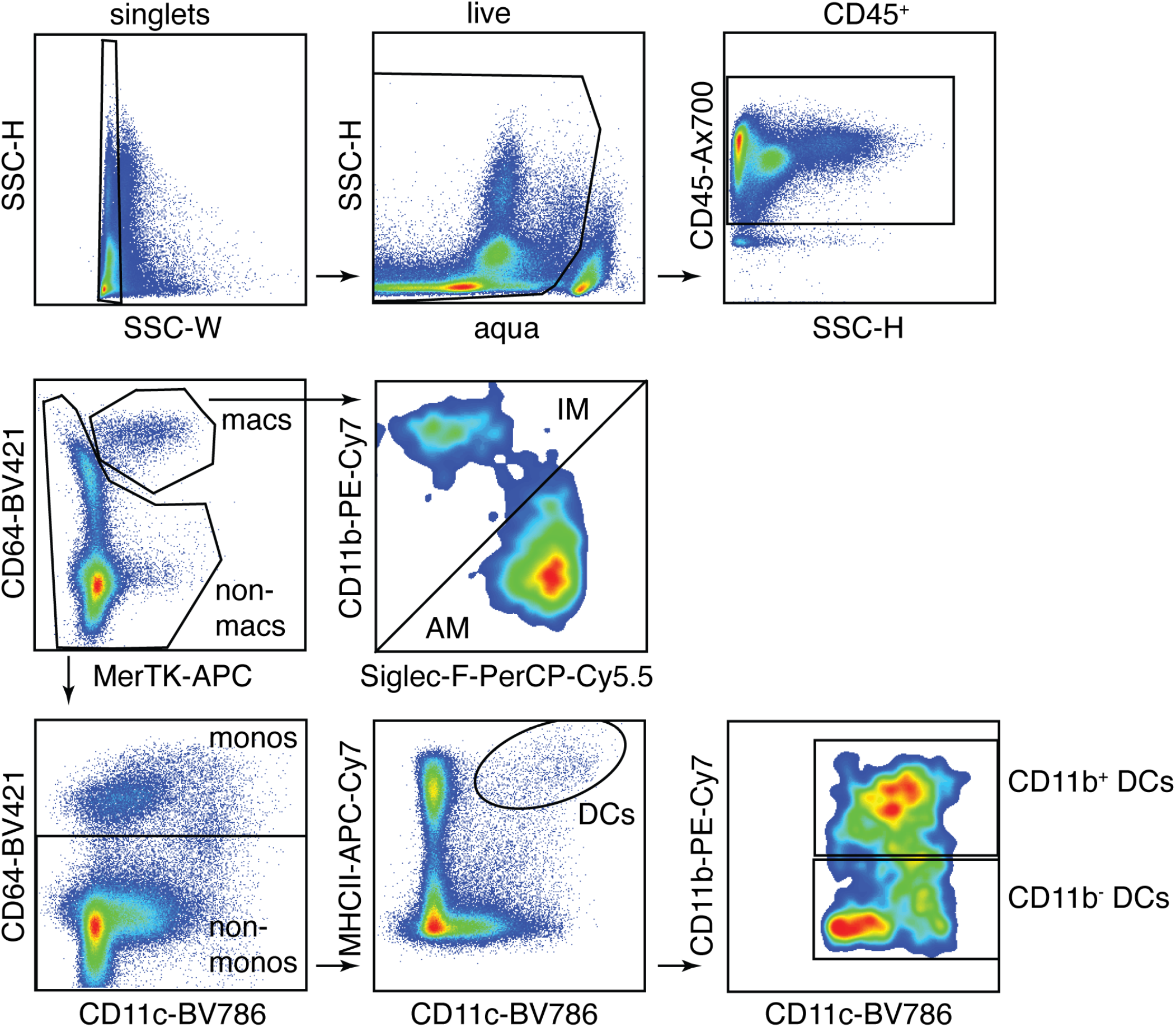
Gating scheme for lung immune cells.

**Supplementary Figure 3.**
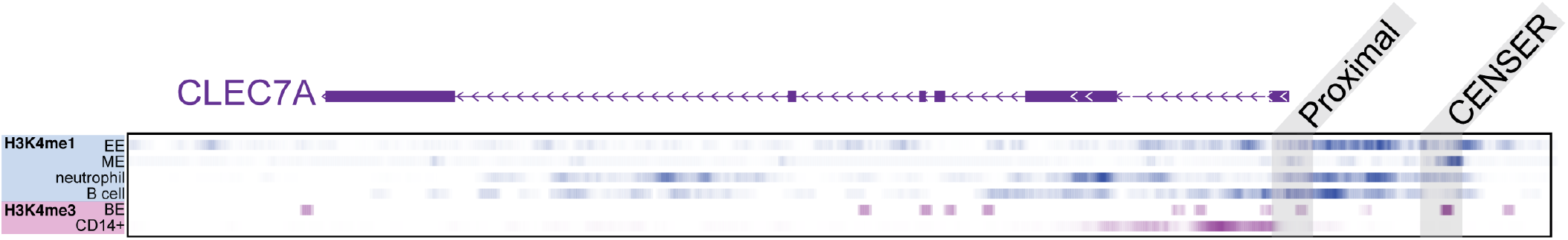
ENCODE ChIP-seq enrichment data for H3K4me1 and H3K4me3 at the human CLEC7A locus. Esophageal epithelium (EE); mammary epithelium (ME), bronchial epithelium (BE).

**Supplementary Figure 4.**
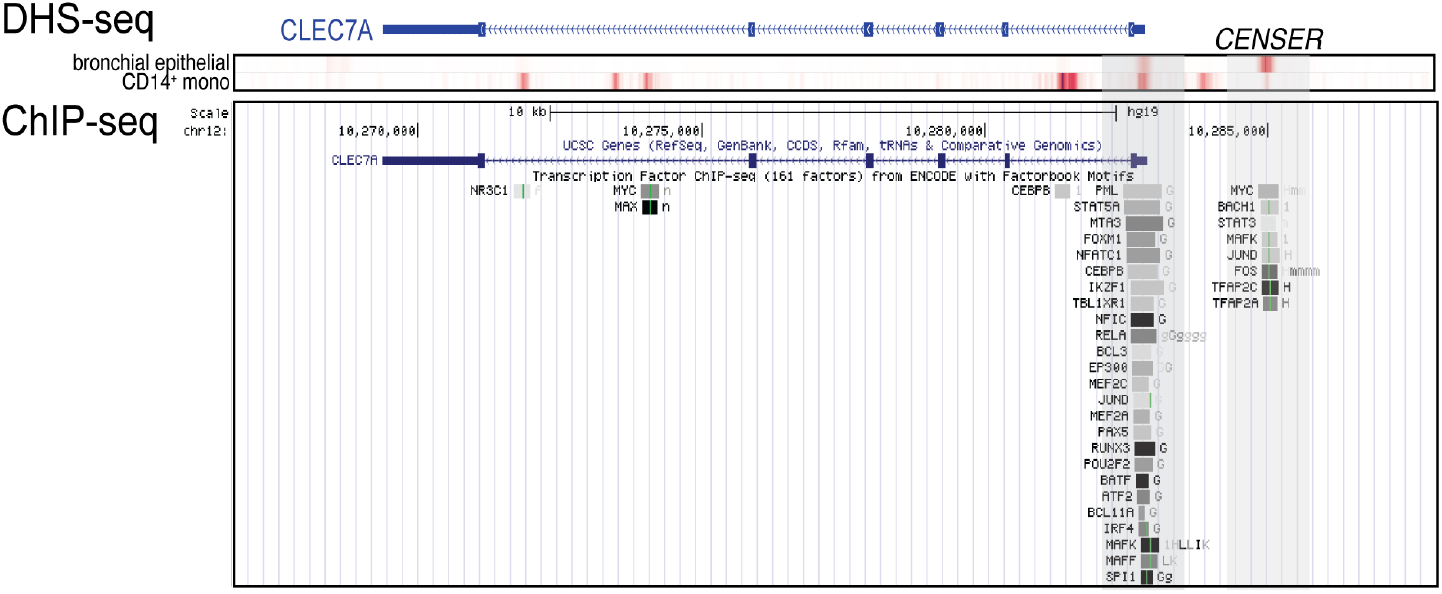
ENCODE DHS-seq and TF ChIP-seq data for the human CLEC7A locus.

**Supplementary Table 1.** List of flow antibodies.

| **Antibody** | **Reactivity** | **Clone** | **Dilution** | **Company** |
| --- | --- | --- | --- | --- |
| CD45-APC-Cy7 | mouse | 30-F11 | 1/800 | BioLegend |
| CD45-Ax700 | mouse | 30-F11 | 1/800 | BioLegend |
| CD31-APC-Cy7 | mouse | 390 | 1/300 | BioLegend |
| CD90-APC-Cy7 | mouse | 30-H12 | 1/200 | BioLegend |
| EpCAM-PE-Cy7 | mouse | G8.8 | 1/600 | BioLegend |
| CD104-APC | mouse | 346-11A | 1/400 | BioLegend |
| CD49f-FITC | mouse | GoH3 | 1/400 | BioLegend |
| Dectin-1-PE | mouse | RH1 | 1/100 | BioLegend |
| CD11c-BV786 | mouse | N418 | 1/400 | BioLegend |
| CD11b-PE-Cy7 | mouse | M1/70 | 1/2500 | BioLegend |
| Siglec-F-PerCP-Cy5.5 | mouse | E50-2440 | 1/400 | BioLegend |
| CD64-BV421 | mouse | X54-5/7.1 | 1/150 | BioLegend |
| MerTK-APC | mouse | 2B10C42 | 1/150 | BioLegend |
| IA-IE-Ax488 | mouse | M5/114.15.2 | 1/3000 | BioLegend |
| CD45-Ax700 | human | 2D1/HI30 | 1/500 | BioLegend |
| CD31-APC-Cy7 | human | WM59 | 1/500 | BioLegend |
| CD90-APC-Cy7 | human | 5E10 | 1/500 | BioLegend |
| EpCAM-PerCP-Cy5.5 | human | 9C4 | 1/250 | BioLegend |
| EpCAM-APC | human | 9C4 | 1/250 | BioLegend |
| CD104-BV786 | human | 439 9B | 1/250 | BD |
| Dectin-1-PE | human | 15E2 | 1/100 | BioLegend |
| STAT3 pTyr705-Ax647 | human | 13A3-1 | 1/100 | BioLegend |
| Zombie Aqua viability dye | all |  | 1/1000 | Biolegend |

**Supplementary Table 2.** Summary of CRS patient characteristics used.

| **Controls** |  |  |
| --- | --- | --- |
| **Age** | **Sex** | **Nasal Polyps** |
| 69 | M | No |
| 63 | F | No |
| 59 | F | No |
| 24 | M | No |
| 50 | M | No |
| 20 | M | No |
| 69 | M | No |
| 43 | F | No |
| 46 | F | No |
| 67 | F | No |
| 50 | M | No |
| **CRSwNP** |  |  |
| **Age** | **Sex** | **Nasal Polyps** |
| 46 | M | Yes |
| 63 | M | Yes |
| 68 | F | Yes |
| 49 | M | Yes |
| 31 | M | Yes |
| 47 | M | Yes |
| 54 | M | Yes |
| 56 | F | Yes |
| 57 | M | Yes |

**Supplementary Table 3.**
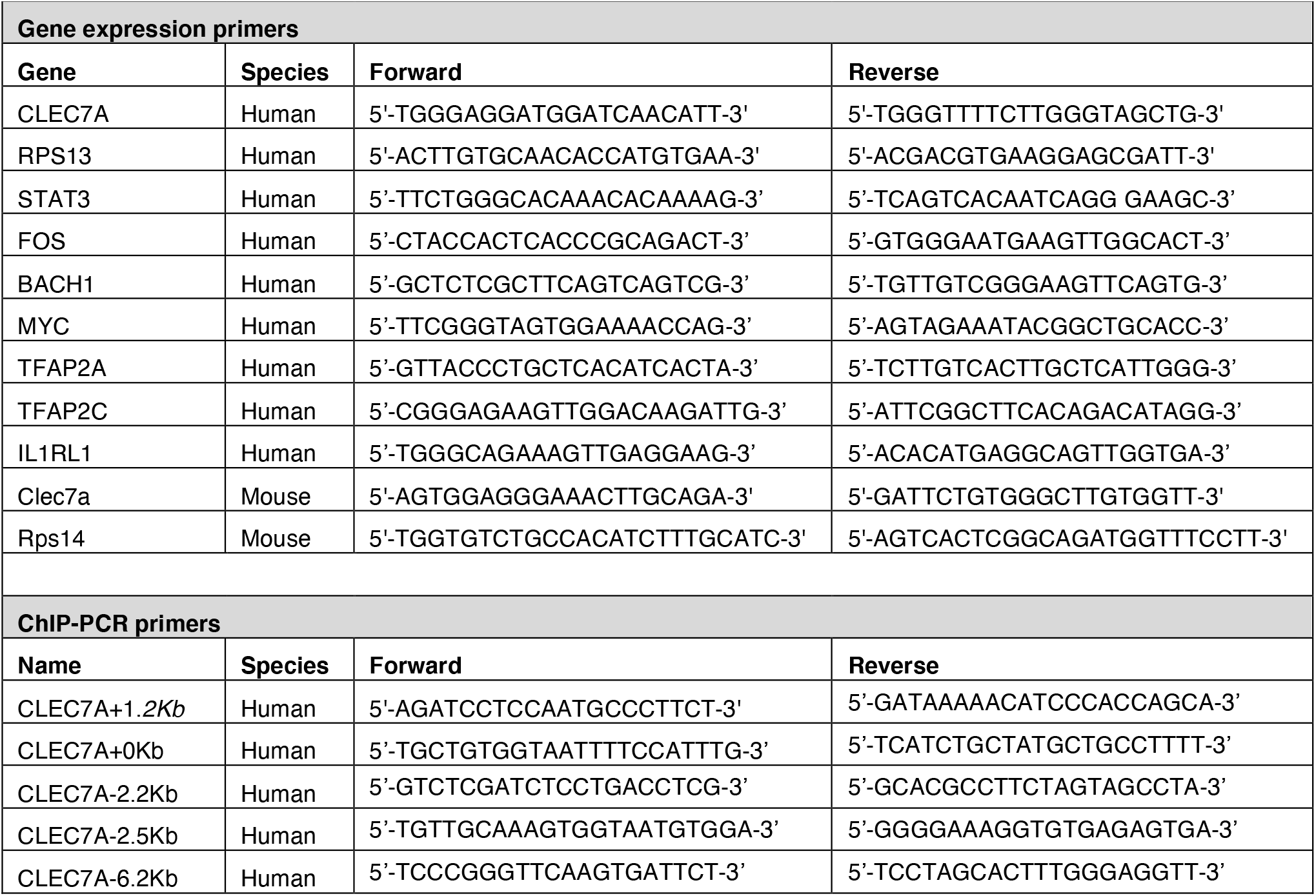
Primer Sequences.

## Notes

### Competing Interest Statement

The authors have declared no competing interest.

